# Key residue on cytoplasmic dynein for asymmetric unbinding and unidirectional movement along microtubule

**DOI:** 10.1101/2022.03.27.485981

**Authors:** Shintaroh Kubo, Tomohiro Shima, Takahide Kon, Shoji Takada

## Abstract

Cytoplasmic dynein 1 is almost exclusively responsible for intracellular transport toward the minus-end of microtubules in animal cells. One of the key factors for the unidirectional movement of dynein is the asymmetry of the unbinding of the motor from the microtubule when an external load is applied; it dissociates more easily from microtubules with minus-end directed loading than with plus-end directed loading. To elucidate the molecular basis for this property, we performed molecular dynamics simulations to identify the key residues responsible for asymmetry, which were then examined experimentally. First, we reproduced asymmetry in the unbinding behavior of dynein using coarse-grained simulations. Then, data analysis together with mutational analysis *in silico* predicted the specific residues that may be responsible for the asymmetry in unbinding. To examine this prediction, we expressed and purified recombinant dynein with mutations in either of the identified key residues. Consistent with the simulations, one of the mutants did not exhibit asymmetry in the *in vitro* unbinding assay. Moreover, the mutant dynein was able to bind and move diffusely along a microtubule but was unable to restrict its movement to the minus-end direction. Our results demonstrate both experimentally and theoretically how the key residue on the microtubule-binding domain generates asymmetry in unbinding, which is a critical mechanism for the unidirectional movement of dynein along a microtubule track.

**Significance Statement:** Cytoplasmic dynein moves to the minus end of microtubules. This unidirectional dynein motility provides the driving force for various cellular activities including vesicle transport, organelle positioning and cell division. One of the key factors for dynein to exhibit unidirectional movement is the asymmetry of unbinding of dynein from the microtubule depending on the direction of external load. By combining computational simulations and *in vitro* experiments, we identified a residue responsible for the asymmetry. A point mutation at the residue indeed abolished unidirectional motility, highlighting the importance of the asymmetric unbinding property in dynein’s unidirectional movement.

## Introduction

Cytoplasmic dynein 1 (dynein) is a two-headed molecular motor that moves to the minus-end of a microtubule (MT). Because dynein drives most of the minus-end-directed cargo transport in animal cells, it plays crucial roles in many cellular processes, such as organelle and nuclear positioning, virus infection, and cell division. Despite the strict directionality of its net movement, each dynein molecule frequently steps towards sideways and backward on a microtubule (1–4) (DeWitt 2012, Qiu 2012, Can 2014, Ando 2020). In addition, two heads in a single dynein molecule demonstrate uncoordinated movement (1, 2, 4) (DeWitt 2012, Qiu 2012, Ando 2020). These unusual features of dynein among cytoskeletal motors have raised the fundamental question of how it achieves net unidirectional movement.

Currently, two major factors are believed to play important roles in the directional control of dynein movement. The first is the direction of the force-generating power stroke. The dynein head consists mainly of ATP-hydrolyzing ring that contains four ATP/ADP binding sites (5) (Kon 2012). A linker and stalk, two slender structures protruding from the ring, connect the cargo-binding tail domain and the microtubule-binding domain (MTBD) to the ring, respectively. Previous studies have demonstrated that ATPase-driven swing-like conformational changes of the linker against the ring is the main contributor to dynein’s force generation, that is so-called “power stroke” (5–9) (Burgess 2003, Shima 2006, Roberts 2009, Kon 2012, Schmidt 2015). Structural studies of dynein-microtubule complexes suggest that the direction of dynein’s power stroke is limited towards the minus-end of the microtubule due to the fixed orientation and tilted angle of dynein against the microtubule (10, 11) (Carter 2008, Imai 2015). Indeed, a protein-engineering study has shown that the net directionality of dynein motility can be altered by changing the orientation of the linker against the microtubule (12) (Can 2019), suggesting that the direction of the power stroke determines the directionality of dynein towards the minus-end of the microtubule.

Another important factor that can determine the direction of dynein movement is the dissociation properties of dynein from microtubules (13–17) (Gennerich 2007, Cleary 2014, Nicholas 2015, Furuta 2017, Ezber 2020). Biophysical studies using optical tweezers have shown that dynein molecules dissociate faster when pulled in the minus-end direction of the microtubule than when pulled in the plus-end direction, which is most likely regulated by the ATPase cycle in the ring domain (14, 15, 17–19) (Cleary 2014, DeWitt 2015, Nicholas 2015, Rao 2019, Ezber 2020). Based on this asymmetry, it has been assumed that intramolecular tension between the two heads of dynein promotes dissociation of the rear head rather than the leading head when both heads bind to a microtubule, biasing the dynein molecule to move forward. Given that the two heads of dynein show apparently uncoordinated motion, the forward stepping bias produced by asymmetric unbinding may be critically important for the processive unidirectional motility of a single dynein molecule.

However, the molecular basis for asymmetric unbinding and how this asymmetry is involved in unidirectional dynein motility remain to be elucidated. In this study, to explore the crucial residues in MTBD for this property, we simulated the behavior of MTBD on a microtubule after applying loads in the plus- and minus-end directions and identified prominent candidates of such residues. Among the candidates, a point mutation on the R3423 residue in MTBD abolished unidirectional dynein movement in both single molecule and ensemble motility assays, although it retained the ability to bind and fluctuate along microtubules. In agreement with the simulation results, this mutant lost the directional dependence of the dissociation rate from the microtubules. These results demonstrate the importance of R3423 in the asymmetric unbinding and unidirectional motility of dynein.

## Results

### Simulating asymmetric unbinding of MTBD

First, we constructed a simulation setup to analyze the directional control of dynein movements (Fig. 1A). This setup contains a *Dictyostelium* dynein MTBD with a proximal portion of the stalk coiled coil and a microtubule (MT) protofilament consisting of three α- and β-tubulin dimers, where the MTBD is strongly bound to the central tubulin dimer in the MT protofilament. Coarse-grained MD simulation of the protein complex was performed using the CafeMol program (30) (Kenzaki et al., JCTC 2011). To quantify the motion of the MTBD, its position, tilt angle, and hinge angle were defined and recorded as follows (Fig. 1B): the position is the center of mass of the globular domain (MTBD) composed of residues A3372 to S3488; the tilt angle *θ* represents the angle between the stalk and MT long axis; the hinge angle of MTBD is the angle formed by the center of MTBD and both ends of the stalk fragment. We also defined the dissociation of MTBD from MT as when the position of the MTBD moved by more than 1 nm from the initial position in our simulation.

**Fig. 1.**
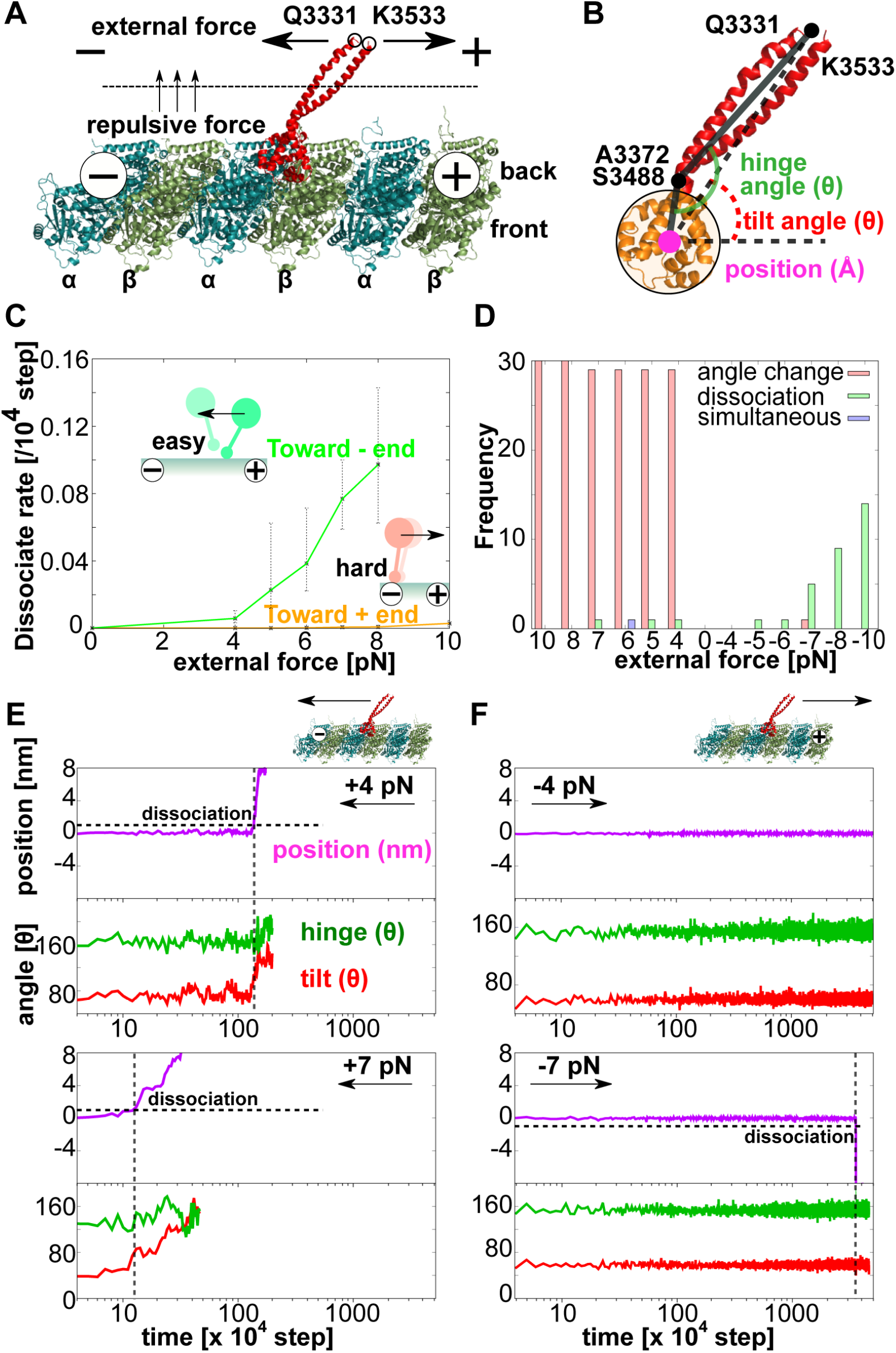
The MD simulations reproduced asymmetric unbinding property of dynein. **(A) The simulation setup.** The MT model used for the simulation was composed of three α-(blue) and β-tubulin (green) dimers in a single protofilament. The MTBD with the short stalk fragments (red) was placed in contact with the gap between α- and β-tubulin in the middle. The MT in this figure is placed with its minus-end on the left. **(B) Definitions of the position, hinge, and MTBD angles in the simulation**. The centroid of MTBD (A3372-S3488, orange) was defined as the position of MTBD (magenta dot). Since A3372 and S3488 is in the hinge region that connect the MTBD to the stalk, we defined the hinge angle (green) as the angle between three points: the midpoint of Q3331 and K3533 at the stalk end, the midpoint of A3372 and S3488, and the MTBD centroid. The MTBD angle means the angle between the line passing through the MTBD centroid and the midpoint of Q3331 and K3533 with the line parallel to the MT long axis. **(C) The dissociation rate under external force**. The curves for the plus-(orange) and minus-end (green) directed loadings demonstrates asymmetry of the dissociation rates. The plots and error bars represent the mean values and standard deviations (SD), respectively; *N* = 30 for each loading condition. **(D) Histograms of the first reaction of MTBD against loadings**. Thirty simulations were carried out for each loading condition, and we counted the frequency of the trajectories with the MTBD angle change prior the dissociation (magenta), the MTBD dissociation without the angle change (green) and simultaneous MTBD angle change and dissociation (purple). **(E, F) Representative trajectories of MTBD movement under loadings**. The trajectories of the MTBD position (magenta), the MTBD angle (red), and the hinge angle (green) with the minus-(E) and plus-end directed loadings (F) are shown on a log time scale. The axes of time and position are plotted on log scales. The upper two panels and bottom two panels show the trajectories with 4 pN and 7 pN of the loadings, respectively.

We then examined whether our computational model could reproduce asymmetric unbinding properties. For this purpose, we simulated the MTBD motion on the MT with and without applying an external force, where 4-8 pN of external forces towards the plus- or minus-end of the MT was applied at the tip of the stalk. When MTBD was pulled toward the minus-end direction, the dissociation rate of MTBD from MT increased rapidly with increasing external forces (Fig. 1C). For example, an external force of 8 pN towards the minus-end direction increased the median dissociation rate up to ∼0.1/10^4^ MD steps. On the other hand, when the same force was applied to the plus-end direction, the dissociation rate remained below 0.01/10^4^ MD steps, showing the clear force direction dependency of the dissociation rate. Therefore, the results validate that our simulation can reproduce the asymmetric unbinding properties of MTBD.

To elucidate the molecular basis of asymmetric unbinding, we further analyzed the behavior of simulated dynein molecules under external forces. The major load-direction-dependent difference in dynein behavior was the stalk angle (Fig. 1D). When the MTBD was pulled in the minus-end direction, the stalk of almost all simulated dynein molecules (176 of 180 trajectories) leaned towards the minus-end, resulting in an increase in the stalk angle against the MT lattice before detachment (Fig. 1D, E). On the other hand, under plus-end directed loading, very few dynein molecules showed significant changes in its stalk angle during its stay on MT (Fig. 1D, F). The stalk angle can be changed by the rotation of the MTBD against the MT and/or stalk bending at the hinge region between the stalk coiled coil and the MTBD. However, regardless of the loading direction, the hinge angle did not change significantly before dissociation of the MTBD (Fig. 1E, F). This result suggests that the rotation of the MTBD, rather than the stalk bending, mainly increases the stalk angle under the minus-end directed loading.

### Identification of the residues responsible for the asymmetry

The loading-direction-dependent MTBD rotation should change the distance between residues involved in MTBD-MT interactions. Because such structural changes in the interaction sites are expected to play a key role in producing the asymmetric unbinding property, we searched for the high contact residues in the dynein-MT complex immediately before its detachment under the minus-end-directed external loading of 4 pN. Using the coarse-grained MD simulation trajectories described above, the probability that each residue pair between MTBD and MT was within 1 nm was calculated. This simulation should reflect all the electrostatic, hydrophobic, and weak interactions in the MTBD-MT complex, because in the coarse-grained force field, weak interaction is incorporated as a force that restabilizes the reference structure. The resulting contact map showed three major interaction regions between the MTBD and MT (Fig. 2A): (1) the MTBD H1-H2 loop with α-tubulin H11’-H12 loop, (2) the MTBD H6-CC2 loop with α-tubulin H11’-H12 loop, and (3) MTBD H3 with β-tubulin H4.

**Fig. 2.**
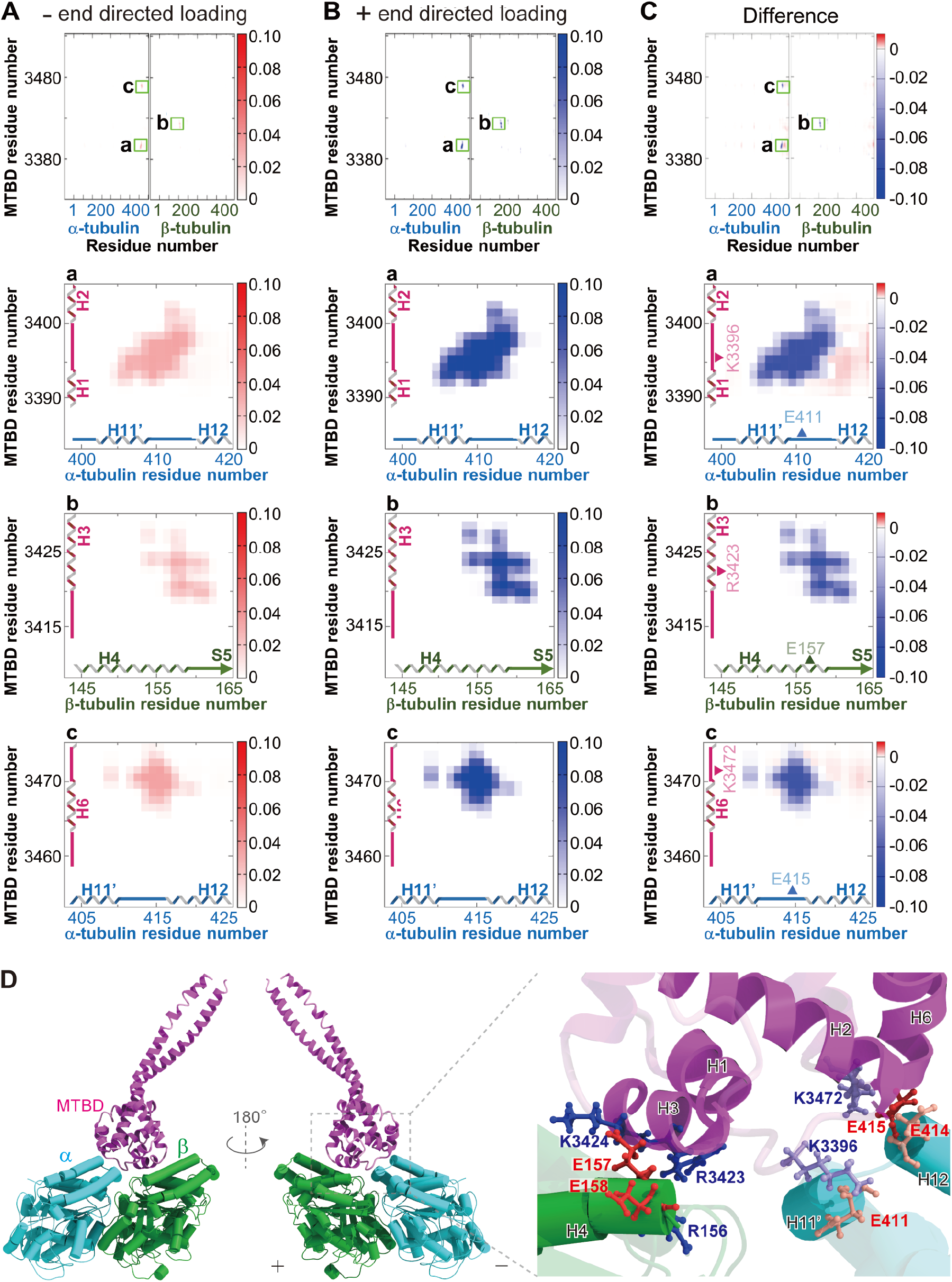
Interaction of MTBD H3 and β-tubulin H4 shows the highest loading directional dependency in the contact probability. **(A-C) Heat maps of the contact probability.** The mean contact probabilities of each residue in MTBD and tubulin dimer under 4 pN of the minus- (A) and plus- (B) end directed loadings and the subtraction of the latter from the former (C) are shown as heat maps. Color scales are on the right of each map. In the contact maps of whole MTBD and tubulin dimer (top panels), the three highly contacted regions are indicated as a-c. The bottom three panels show the enlarged views of these three regions. In the light-green boxes named a, b, and c are the highest contact area picked up the right side. **(D) Detailed structure of the MTBD-MT interface**. MTBD, α-, and β-tubulin are colored magenta, blue, and green, respectively. In the domains with the high contact probabilities (opaque ribbons and cartoons in the right panel), positively-charged (Arg and Lys) and negatively-charged residues (Glu) are indicated as blue and red balls and sticks, respectively.

To evaluate the loading-direction dependency of these contact regions, we calculated the same contact map in the case of plus-end directed loading of 4 pN. As a result, the same three regions, as in the case of minus-end loading, were shown to be the major contact regions (Fig. 2B). However, the loading directional dependency of the contact probability differed among the regions. The MTBD H3-β-tubulin H4 region demonstrated the largest difference in contact probability, depending on the loading direction (Fig. 2C). Among the top 20 pairs that showed a large loading directional dependency in contact probability, 17 pairs located in this region (Table S1). In comparison to the minus-end-directed loading, the plus-end-directed loading led to a higher contact probability in this region, suggesting that the MTBD-MT interaction in this region is stronger under plus-end-directed loading. The other two regions (the MTBD H1-H2 loop-α-tubulin H11’-H12 loop and the MTBD H6-CC2 loop-α-tubulin H11’-H12 loop) also showed higher contact probabilities under plus-end-directed loading than under minus-end-directed loading, likely contributing to the asymmetric unbinding. However, these MTBD loops interact with the α-tubulin C-terminal H12 only when a force is applied in the minus direction, compensating for the decreased interactions with the α-tubulin H11’-H12 loop (Fig. 2C). Therefore, the net contact probability of these two MTBD loops with MT showed less loading-directional dependency than the MTBD H3-β-tubulin H4 region, underscoring the importance of the MTBD H3-β-tubulin H4 region in the asymmetric unbinding property. Considering the contact probability and electrostatic effects, we identified that the major contributors to the MTBD H3-β-tubulin H4 contact should be R3423/K3424 in the MTBD H3 domain and R156/E157/E158 in the β-tubulin H4 domain (Fig. 2D). Among these charged residues, E158 in β-tubulin was found to orient the opposite side towards R3423/K3424 in MTBD. This suggests that the interactions between R3423/K3424 (MTBD H3) and R156/E157 (β-tubulin H4) are responsible for the asymmetric unbinding of dynein.

We further performed *in silico* mutant analysis to investigate the effects of the above mentioned three MTBD regions on the asymmetric unbinding. Same as R3423 and K3424 in H3, K3396 in the H1-H2 loop and K3472 in the H6-CC2 loop are considered to be responsible residue for the interaction between the loops with tubulin, based on electrostatic interactions and structural information. Therefore, we simulated the dissociation rate of the dynein under loading when these four basic residues including R3423 and K3424 were mutated to neutral alanine or acid aspartate. As a result, in the case of K3396 mutation, K3396A demonstrated the highest dissociation rates under plus-end directed loading, while the wild-type rarely dissociated under the same condition (Fig. S1A). Contrarily, for R3423, R3423D showed a higher dissociation rate than R3423A under plus-end directed loading (Fig. S1B). Similar to the case of R3423, K3424D showed the highest dissociation rate among K3424 mutants, but this mutant showed a very low affinity for microtubules and rapid dissociation from microtubules even under no-loading conditions (Fig. S1C). On the other hand, K3424A retained intact level of affinity to microtubule while it increased the dissociation rate under plus-end directed loading. For K3472, neither of the mutations increased the dissociation rate under plus-end directed loading (Fig. S1 D). Considering the results of all these MD simulations, we decided to use K3396A, R3423D and K3424A mutants in the following *in vitro* experiments to verify the effects of the residues in the asymmetry.

### *In vitro* experiments to examine the role of the key residues on dynein motility

To evaluate the role of MTBD R3423/K3424 residues on dynein motility, we performed *in vitro* motility experiments using mutants of the head of *Dictyostelium* cytoplasmic dynein heavy chain. First, motile activity was tested by microtubule-gliding assay, in which microtubules were translocated by multiple dynein molecules attached on the glass surface (Fig. 3). As shown previously (Kon et al. 2004, Shima et al. 2006), the wild-type head robustly drove gliding movement of microtubules (Fig. 3A, 3B) with the mean velocity of 1.82 ± 0.29 μm s^-1^ (mean ± standard deviation (SD), n = 150) in the presence of 1 mM ATP. On the other hand, R3423D induced only extremely slow microtubule movement; the mean velocity of microtubule gliding on the R3423D coated surface was 0.02 ± 0.02 μm s^-1^ (mean ± SD, n = 116), which is approximately 100 times slower than that of the wild-type (Fig. 3C). Detailed kymograph analysis demonstrated that the wild-type drove unidirectional microtubule movement, whereas R3423D mainly showed slow one-dimensional fluctuation of microtubules (Fig. 3D). This severe effect of the mutation on microtubule gliding motility is consistent with the above-mentioned prediction using *in silico* simulations. In contrast, the other two mutants, K3396A and K3424A, drove robust unidirectional microtubule movement similar to the wild-type (Fig. 3E-H), highlighting the critical importance of R3423 among residues in the MTBD on dynein’s unidirectional motile activity.

**Fig. 3.**
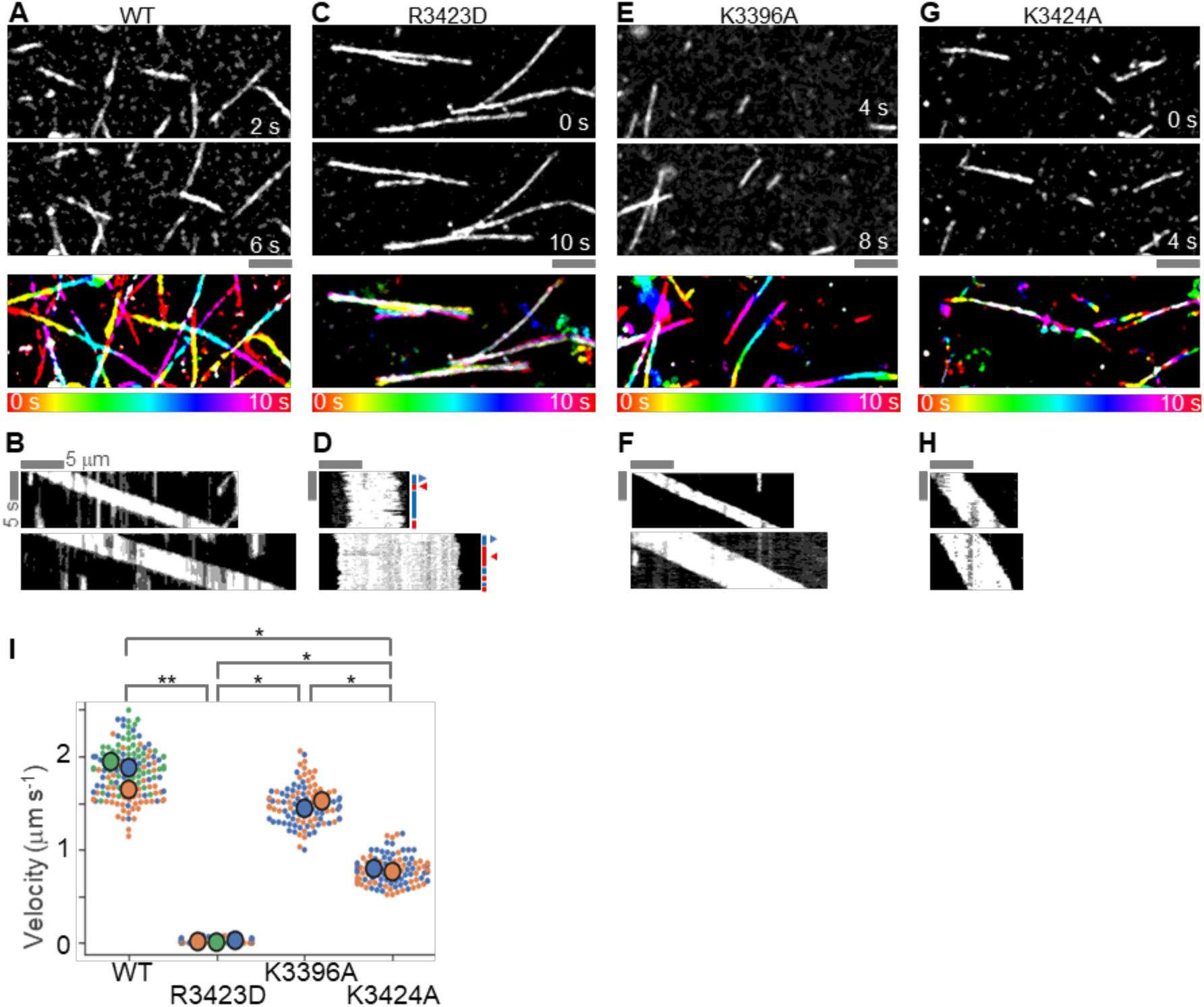
R3423D mutation hampered unidirectional microtubule gliding motility of dynein head. **(A-H)** Dark field microscopic images and kymographs of microtubules on the dynein-coated surface (**A, B**, wild-type; **C, D**, R3423D; **E**,**F**, K3396A; **G**,**H**, K3424A). (**A, C, E, G**). In addition to the mutations on R3423/K3424 residues, K3396A mutant was also tested as a reference. Time lapse images. The top and middle panels show images of microtubules at the specific time points. The bottom panels show temporal color images. Scale bar, 5 μm. (**B, D, F, H**) Kymographs of the microtubule movement. Direction of microtubule gliding was indicated by blue and red arrow heads in (**D**). (**I**) Dot plots of the microtubule gliding velocities. Each color of the dots represents one of three repetitive experiments. The bigger circles show the mean velocities of each experiment. \**P* < 0.05 and \*\**P* < 0.001 (T-test using the SuperPlot method) (Lord et al. 2020).

A possible cause of this slow and fluctuated microtubule movement by R3423D could be that the mutation suppresses the ATP hydrolysis cycle of dynein and stops the power stroke movement. To test the possibility, we measured ATPase activity of R3423D. Similar to the previously reported ATPase activity of the wild-type monomer (Kon 2004), ATPase activity of R3423D monomer was activated by presence of microtubule [8.6 ± 0.8 s^-1^ (mean ± SD) without microtubule, 14.8 ± 1.9 s^-1^ (mean ± SD) with 10 μM of microtubule; four repetitive assays were performed for each condition]. The results showed that the mutation does not stop the dynein ATP hydrolysis cycle. Also, the ATPase activation by microtubule suggests that R3423D can take a strong binding state to microtubule during its mechanochemical cycle, which is needed to perform the power stroke. This notion was further supported by the single molecule observation of GST-dimers of the dynein head under a total internal reflection fluorescence microscope (TIRFM). Some of the R3423D dimer molecules bound and moved along microtubules (Fig. 4), suggesting that the mutant retained the ability to bind to microtubules. Similar to the case of microtubule-gliding assay, the directionality of movement clearly differed between the wild-type and R3423D mutant dimers. As previously reported (Numata et al., 2011; Imai et al., 2015), the wild-type dimer exhibited overall unidirectional movement towards the minus-end of the microtubule (Fig. 4A). On the other hand, the R3423D dimer frequently switched the direction of the movement (Fig. 4B). From the traces of movement along the microtubules, we evaluated the components of both active motile and one-dimensional diffusive movement. The mean square displacement plots demonstrated that the diffusion constant of the R3423D dimer was seven times higher, while the active motile velocities of R3423D were less than 1/6 of those of the wild-type (Fig. 4C, 4D, Table 1), resulting in the short run length of the movement (Table 1). These results suggest that the R3423D mutant retains microtubule binding ability but loses ability to actively step towards the minus-end direction.

**Fig. 4.**
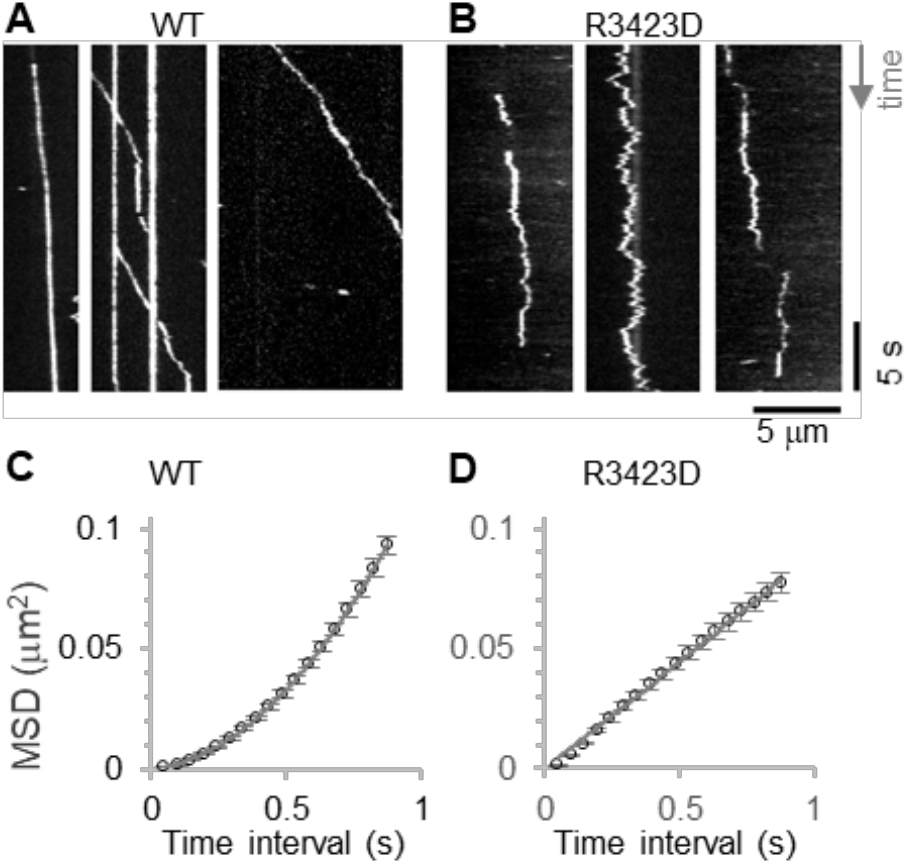
R3423D dimer showed highly fluctuating movements along microtubule. **(A, B)** Kymographs of the dimeric dynein movement along microtubule (**A**, wild-type; **B**, R3423). (**C, D**) Mean square displacement plots (**C**, wild-type; **D**, R3423). By fitting the data with quadratic function, the diffusion constants and active motile velocities were calculated. The values obtained are listed in Table 1.

**Table 1.** Summary of dimeric dynein movement along microtubule. Each value indicates the mean ± standard error of the mean (SEM)

| Construct | Active motile velocity<br>$\mu\text{m s}^{-1}$ | Diffusion constant<br>$\times 10^{-3} \mu\text{m}^2 \text{s}^{-1}$ | Attachment rate<br>$\mu\text{m}^{-1} \text{nM}^{-1} \text{s}^{-1}$ | Duration<br>s | Run length<br>$\mu\text{m}$ | N |
| --- | --- | --- | --- | --- | --- | --- |
| WT | $0.33 \pm 0.07$ | $5.9 \pm 1.7$ | $0.06 \pm 0.03$ | $1.2 \pm 0.7$ | $0.8 \pm 0.1$ | 51 |
| R3423D | $0.05 \pm 0.08$ | $43 \pm 2$ | $0.02 \pm 0.02$ | $0.8 \pm 0.6$ | $0.2 \pm 0.1$ | 47 |

Yet, the lower attachment rate and shorter duration of the mutant molecule on microtubules in the single-molecule motility assay (Table 1) indicate that the mutation decreases the binding affinity of dynein for microtubules. It is possible for a single GST-dimer molecule to fluctuate along microtubules only due to low binding affinity, even if the asymmetric unbinding property remains. For example, mammalian cytoplasmic dynein 1 molecules in the autoinhibited state have shown similar diffusive movements along microtubules (Torisawa et al., 2014; Zhang et al., 2017). However, in the microtubule-gliding assay system, a large number of dynein molecules interact with a single microtubule (Shima et al., JSB, 2006), and therefore, the low binding affinity alone cannot explain the fluctuating movement of the microtubules on the R3423D-attached surface. When power stroke of a dynein molecule causes physical tension between multiple dynein molecules bound to a same microtubule, asymmetry in unbinding makes the molecules pulling the microtubule backwards to dissociate from the microtubule, resulting in the unidirectional microtubule movement. Indeed, the aforementioned mammalian cytoplasmic dynein also has shown unidirectional movement when multiple dynein molecules were linked and simultaneously interact with one microtubule (Torisawa et al., 2014). Therefore, the result that microtubules showed fluctuating motion despite the binding of many R3423D molecules in the microtubule-gliding assay (Fig. 3D) supports the notion that the mutant lost the asymmetry in unbinding, as expected from the MD simulations.

To evaluate the asymmetric unbinding property of R3423D, we then applied hydrodynamic forces to dynein-coated polystyrene beads on microtubules by shear flow using a microfluidic system (42) (Urbanska et al., 2021). Consistent with the notion, the shear flow needed to unbind R3423D-coated beads from microtubules did not show directional dependence, while the wild-type coated beads required twice faster flow when pulled towards the plus-end of the microtubule compared to when pulled to the minus-end (Fig. S2). Together, these results suggest that the R3423 residue is responsible for the asymmetric unbinding of dynein, and that the asymmetric unbinding property is critical for the unidirectional motility of dynein along a microtubule track.

## Discussion

Our *in silico* and *in vitro* experiments highlighted the importance of the R3423 residue in the asymmetric unbinding property and unidirectional movement of dynein. The MD simulations of dynein behavior under load clearly showed that the strength of the interactions between R3423/K3424 in MTBD and E157/E158 in β-tubulin varied with the direction of the applied load. In line with the hypothesis proposed by the simulations, our *in vitro* experiments showed that the R3423D mutation abolished the loading direction dependence of the unbinding rate and disrupted the ability of dynein to move unidirectionally along the microtubules. Because the main role of cytoplasmic dynein is the directional transport of various cargoes toward the minus-ends of microtubules (43, 44) (Paschal & Vallee, 1987, Reck-Peterson, 2018), this key residue for asymmetric unbinding is also critical for cellular dynein function.

The molecular basis of how R3423 contributes to the asymmetric unbinding property can be explained by the electrostatic interaction between MTBD and β-tubulin. As suggested by the contact map (Fig. 2), R3423 and K3424 of MTBD are positioned to sandwich E157 of β-tubulin, where R3423/K3424 forms electrostatic interactions with E157 (Fig. 5A, B). On the other hand, R3423 is also in close proximity to R156 of β-tubulin, where the two residues are electrostatically repulsive, suggesting that these electrostatic attraction and repulsion forces are balanced to form the MTBD-MT interaction under no-load conditions (Fig. 5B). Under plus-end directed loading, the simulation shows that the MTBD rotates so that the stalk-MTBD junction translocates toward the plus-end, pushing R3423/K3424 closer to E157 and away from R156 (Fig. 5C), resulting in the enhancement of the MTBD-β-tubulin interaction. In contrast, minus-end directed loading rotates and shifts MTBD against the microtubule lattice in the opposite direction, pushing R3423/K3424 away from E157 and closer to R156 in β-tubulin, which could weaken the interaction (Fig. 5C). Collectively, the asymmetric unbinding property could arise from changes in the strength of the electrostatic interaction between MTBD R3423/K3424 and β-tubulin R156/E157 depending on the direction of the applied load. With regards to why the K3424A mutation abolished the asymmetry in the unbinding and unidirectional motility of dynein, our simulations show that even in the K3424A mutant, the R3423-E157 interaction has a strong loading-direction dependence, becoming stronger with loading in the plus-end direction and weaker with loading in the minus-end direction (Fig. 5D). In contrast, in the R3423D mutant, the loading direction dependence was lost owing to the switching of salt bridges, and the K3424-E157 interaction became weaker with the minus-end directed loading compared to that with the plus-end directed loading. However, the R3423D-R156 salt bridge was newly created to compensate for the loss of the K3424-E157 salt bridge. As a result, the overall interaction strength between MTBD and β-tubulin did not change markedly (Fig. 5E). Therefore, the formation of the salt bridge with R156 is likely to be responsible for the different effects of the R3423D and K3424A mutations on the asymmetric unbinding property.

**Fig. 5.**
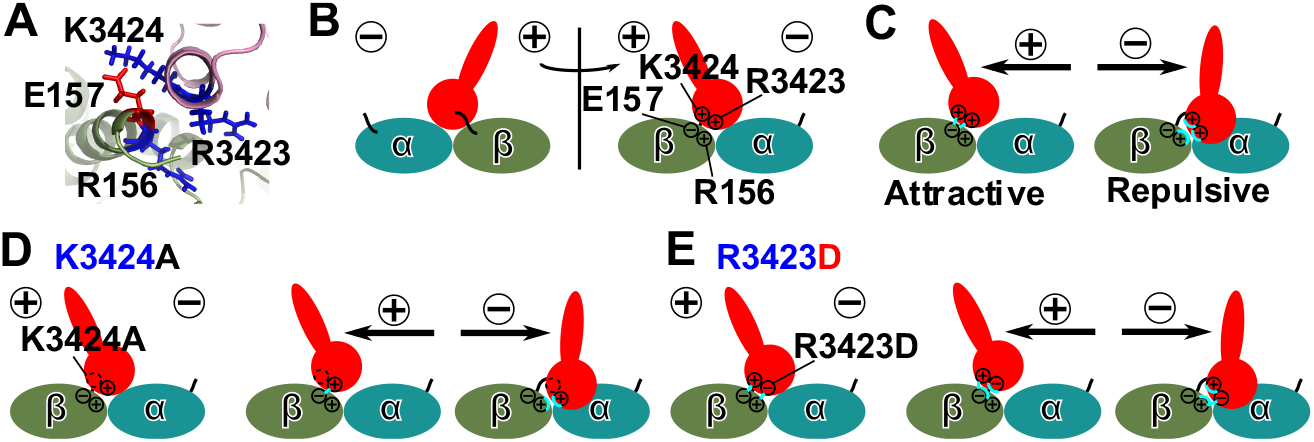
The electrostatic effects on R3423 can explain the asymmetric unbinding property of dynein. **(A, B)** Structural configuration of the key residues. **(A)** 3D model. **(B)** Schematic drawings. **(C-E)** Schematic drawings of the electrostatic effects on MTBD R3423/K3424 with <u>β</u>-tubulin R156/E157 under the external loadings. In the case of wild-type **<u>(C)</u>**, K3424A **(D)**, and R3423D **(E)** mutants are shown. **(D, E)** The left, middle, and right panels shows in the cases of no-, the plus-end directed, and minus-end directed loadings, respectively.

The other contact sites on the microtubules are mainly located in the C-terminal region of α-tubulin. Our simulations showed that although load-dependent salt-bridge switching occurs in these interaction regions, the overall strength of the interaction between MTBD and α-tubulin did not change significantly. Under plus-end directed loading, the H11’-H12 loop in α-tubulin formed salt bridges with the H1-H2 and H6-CC2 loops of MTBD. Under minus-end directed loading, the loops of MTBD were shifted away from the H11’-H12 loop and bound with neaby residues in H12 of α-tubulin instead (Fig. 2C). Since the H11’-H12 loop and H12 of α-tubulin are both negatively charged and highly mobile regions suitable for accommodating the interaction with the positively charged H1-H2/ H6-CC2 loops of MTBD, the MTBD-α-tubulin interaction can be maintained even during the loading-dependent conformational change of MTBD. Although these regions are unlikely to be responsible for asymmetric unbinding, they play important roles in dynein motility (10, 20, 45) (Carter 2008, RedWine 2012, McKenney 2016). The C-terminal region of α-tubulin, including the H11’-H12 loop, is known to be a major binding site for MAPs (46, 47) (Roll-Mecak 2015, Volkov 2020), which may affect the motility of dynein. In addition, a previous study suggested that the H11’-H12 loop of α-tubulin is an important region for dynein to exert its motor activity and is essential for the microtubule-dependent stimulation of dynein ATPase (48) (Uchimura 2015). Consistent with this report, the present study has shown that MT-stimulated ATPase activity is maintained even in the R3423D mutant, suggesting that the two mechanisms essential for dynein to exert its motor activity, namely the microtubule-dependent activation of ATPase and the asymmetric unbinding properties of MTs, are mutually independent mechanisms that depend on different MTBD-MT interactions.

Our study also highlighted the importance of asymmetric unbinding in the unidirectionality of dynein motility. The R3423D mutation identified and examined in this study not only disrupted the asymmetry of unbinding but also abolished the unidirectionality of dynein motility. Herein, we discuss how asymmetric unbinding contributes to the unidirectionality of dynein motility. We assumed that a dynein molecule with two heads stepped along a microtubule with a large step size. Under these conditions, intramolecular tension between the two heads in dynein leads to dissociation of the rear head from the microtubule, causing the entire dynein molecule to move forward, as shown in previous single-molecule studies (1, 2) (Qiu 2012, DeWitt 2012). The asymmetric unbinding mechanism should be essential for this tension-dependent dissociation of the rear head; if the unbinding rate is independent of the loading direction, as is in R3423D mutant, the front and rear heads can randomly dissociate from the microtubule, making the dynein molecule take forward and back steps evenly. Even when the dynein molecule steps with a small step size that does not generate intramolecular tension between the heads, the asymmetry of unbinding remains important. Since dynein molecules moving in the minus-end direction of the microtubule are always subjected to the drag force in the plus-end direction from the load (49) (Palenzuela 2020), it would be necessary for dynein to have an MTBD that has a high binding affinity for MT, especially against the plus-end directed forces. Thus, in addition to the direction of the power stroke, the asymmetry of unbinding from microtubules is an important basis for the unidirectionality of dynein motility. Our work, together with further studies on the asymmetry of dynein in the weak binding state, will contribute to a complete understanding of the relationship between dynein’s mechano-chemical cycle and motility.

## Methods

### Protein models

We used *Dictyostelium discoideum* cytoplasmic dynein-1, of which sequence was obtained from UniProtKB (P34036). In the modeling and analysis, we used only Q3331-K3533, which contains full of the MTBD and a part of the stalk. The reference structure of the high-affinity MTBD was modeled by combining the two model structures in the following protocol. The first model is based on pdb ID: 3J1T (20) (Redwine Science 2012), which contains structures of the MTBD and short helical turns in the stalk for *Mus musculus*, and is based on cryo-electron microscopy data in the complex with MT and computational modeling. Using this template, we constructed a homology model. The second model used was pdb ID: 3VKH (5) (Kon Nature 2012), which is the X-ray crystal structure of *Dictyostelium discoideum* cytoplasmic dynein in the ADP-bound form. While this structure contains the MTBD, its resolution near the MTBD is apparently low. Thus, we decided to use this model for a part of the stalk that was not included in the first model. Then, the two models were integrated via molecular simulation, as previously described (21) (Kubo, PLoS CP). For MT, we used pdb ID: 3J6F (22) (Alushin Cell 2014) as the reference structure. Using the UniProtKB(A0A0M3KKT1) and UniProtKB(A0A0M3KKT2) sequences of *α-*tubulin and *β*-tubulin, respectively, we obtained full-length structures via loop modeling using MODELLER (23) (Šali and Blundell 1993). We then modeled the complex structure of MT and MTBD by superimposing the above-mentioned models on 3J1T. In order to remove locally unfavorable configurations, the energy minimization calculation was performed using the all-atom MD simulation using the GROMACS version 5.1.1 (24, 25) (Abraham SoftwareX 2015, Pronk Bioinfomatics 2013) with the amber99sb-ildn force field for proteins (26) (Lindorff-Larsen 2010) and the TIP3P model for water (27) (Jorgensen 1983). Energy minimization was performed using the steepest descent minimization algorithm.

### Coarse-grained simulation

With the atomic structures of the strong-binding affinity MTBD structures in the dynein head, we performed the coarse-grained MD simulations; the two ends of MTBD (Q3331 and K3533) are pulled by constant force toward the longitude direction of the MT. The amplitude of pulling force was 0, ±4.0 pN, ±5.0 pN, ±6.0 pN, ±7.0 pN or ±8.0 pN. Throughout this study, the total simulation time was 5000 × 10^4^ MD steps. Simulations were performed 30 times for each external force. Here, the plus and minus symbols represent the direction of the pulling force. In reality, the steric hindrance of the ring domain limits the MTBD-stalk angle against MT. To reflect this effect, an additional repulsive force Δ𝔽 was applied to the model MTBD in the orthogonal direction to the MT protofilament as follows:

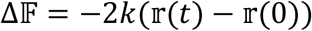

where 2*k* is the spring constant, which is set to *k* =100, and 𝕣(*t*) is the relative position of the two MTBD when the simulation time is *t*. The detachment rate was defined as the reciprocal MD step at which the position of the MTBD moved by 1 nm from the initial position. When the MTBD did not move by 1 nm by the end of a simulation 5000 × 10^4^ MD step, we used the final step for the estimation.

### Atomic interaction-based coarse-grained model (AICG2+)

We used a previously developed and well-tested coarse-grained model AICG2+, in which each amino acid is represented by a single bead located at the Cα position (28, 29) (Li, Terakawa, Wang, and Takada 2012; Li, Wang, and Takada, 2014). While AICG2+ is a structure-based model explicitly biased towards the reference structure, all terms are tuned to represent atomistic chemical interactions in the reference structure. The energy function is expressed as follows:

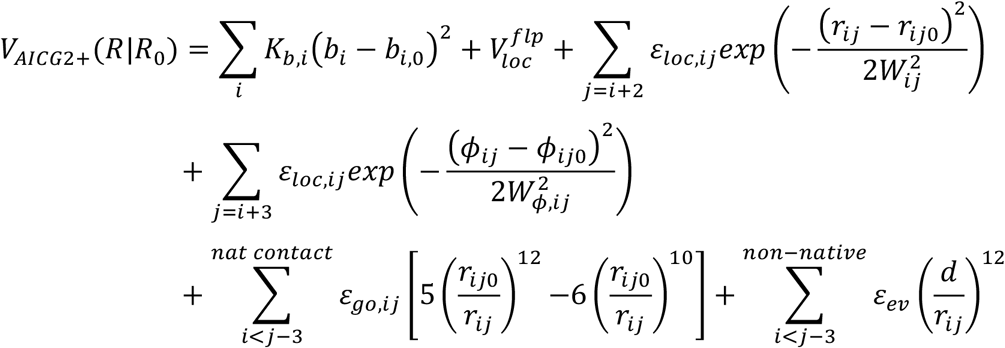

Here, each term represents the elasticity of the virtual bond between two consecutive Cα’s, the sequence-dependent local potential made of virtual-angle- and virtual-dihedral-angle terms, the structure-based local potential between the *i*-th and *i*+2-th residues, the structure-based local potential for dihedral angles, the contact potential for non-local natively interacting pairs, and the generic repulsion for the remaining non-local pairs, in that order. The vector *R* stands the 3*n*_*aa*-_ dimensional Cartesian coordinates of the target protein, where *n*_*aa*_ is the number of amino acids in the protein. *R*_0_ is the corresponding coordinate in the reference structure (all variables with subscript 0 indicate the parameters that take the corresponding values in the reference structure). *b*_*i*_ is the *i*-th virtual bond length between the *i*-th and *i*+1-th amino acids. *r*_*ij*_ is the distance between the *i*-th and *j*-th residue. *ϕ*_*ij*_ is the dihedral angle defined as the *i*-th, *i*+1-th, *i*+2-th, and *i*+3-th residue. 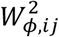 is a parameter that represents the width of the attraction. *K*_*b*_,_*ibd*_, *ε*_*loc,ij*_, *ε*_*ev*_, and *d* are parameters. For the meaning and default values of these parameters, the CafeMol manual can be consulted (30) (Kenzaki et al., 2011)

### Create contact maps

To decipher the mechanistic origin of the anisotropic detachment, we investigated which amino acids of MTBD (A3372-S3488) and MT (entire alpha- and beta-tubulin) make contact when the MTBD is to detach. Each pair of residues in MTBD and MT were assigned to be in contact when they were within 1 nm. To reduce the noise for each pair of residues, we took the average contact probabilities for the last 100 frames (one frame corresponds to10^4^ MD steps) before the detachment for each trajectory. If detachment occurred before the 100× 10^4^*th* MD step, we used all frames before detachment. This was a contact map created from a single trajectory. Because we performed 30 simulations for each setup, the final average contact maps for each setup were an average of the 30 overlaid contact maps.

### Protein preparation

Tubulin was purified from porcine brain through four cycles of polymerization and depolymerization using 1 M PIPES buffer (1 M PIPES-KOH, 1 mM EGTA, and 1 mM MgCl_2_, pH 6.8) to effectively remove microtubule-associated proteins (31). Labeled tubulin was prepared by incubating polymerized microtubules with tetramethylrhodamine succinimidyl ester (C-1171; Thermo Fischer) or NHS-LC-biotin (21336; Thermo Fisher Scientific) for 30 min at 37°C (32). The labeling reaction was quenched by adding 2 mM l-glutamine (16919; Nacalai). Then, we purified the functionally labeled tubulin using two cycles of polymerization and depolymerization.

The expression plasmids for the C-terminal 380-kDa fragment (V1388-I4730) of *Dictyostelium discoideum* cytoplasmic dynein heavy chain fused at its N-terminus with His6-FLAG-biotin-Glutathione S-Transferase (HFBGST380) or His6-FLAG-biotin (HFB380) tandem tags, and its AAA2 with SNAP-tag, have been previously described (33) (Numata et al., 2011). As the biotin tag, we used the C-terminal 72 residues (G524–A595) of the *Klebsiella pneumoniae* oxalacetate decarboxylase α-subunit (Shima et al., 2006). The three stalkhead mutants were created using a QuickChange mutagenesis kit (Agilent), according to the manufacturer’s instructions.

Plasmids were introduced into Dictyostelium cells by electroporation. Transformed cells were selected and harvested as previously described (34) (Kon, Shima, Sutoh, 2009). The expressed dynein heavy chains were purified by nickel-nitrilotriacetic acid and anti-FLAG affinity chromatography (34) (Kon, Shima, Sutoh, 2009). Heavy chains with biotin tags were almost fully biotinylated *in vivo*. The protein concentrations were determined by the Bradford method (35) (Bradford, 1976) using BSA as a standard. A fraction of dynein molecules was labeled with SNAP-surface 649 (New England Biolabs) and purified using micro Bio-spin P30 gel filtration columns (33) (Numata 2011; BioRad).

### Fluorescent microscopy

Non-labeled tubulin, tetramethylrhodamine-labeled tubulin, and biotin-labeled tubulin were copolymerized to yield microtubules with rhodamine and biotin labeling ratios of 2% and 5%, respectively. Tubulin was copolymerized in the presence of 1 mM GTP. The polymerized microtubules were stabilized by adding 0.1 mM paclitaxel (LC Laboratories). The glass chambers used for the observation were prepared as follows: after sonication in 1N KOH and plasma treatment (Diener), the surfaces of the cover glasses (C022221S; Matsunami Glass) were silanized with N-2-(aminoethyl)-3-aminopropyl-triethoxysilane (KBE-603; Shin-Etsu Chemical), followed by incubation with 200 mg/mL NHS-PEG (ME-050-TS, NOF) with and without 1 mg/mL NHS-PEG-biotin (BI-050-TS; NOF) for 3 h at room temperature to make PEG-biotin-coated and PEG-coated glasses (36) (Osuka 2018). The PEG-biotin-coated glass and PEG-coated glass were separated using a 30-μm layer of double-sided tape (5603; Nitto-Denko) to create the flow chamber. The microtubules were immobilized on the PEG-biotin-coated glass surface via neutralized avidin (011-24233; Fuji film-Wako) and then, the glass surface was blocked with imaging solution (12 mM PIPES-KOH, 2 mM MgSO_4_, 1 mM EGTA, 1% (w/v) Pluronic F-127, 1 mg/mL casein, 1 mM d-biotin, 2 mM dithiothreitol, 0.2 mg/mL glucose oxidase, 40 μg/mL catalase, 1 mM glucose, pH 7.0). Finally, fluorescently-labeled dynein was added to the imaging solution and introduced into the chamber. The samples were illuminated with 532-nm or 642-nm lasers (Coherent). The fluorescent images of microtubules and dynein molecules were observed using total internal reflection fluorescence microscopy (Ti2, Nikon) with a 100× objective lens (Apo TIRF, NA 1.49, oil) (Nikon) and recorded at a frame rate of 20 frames per second using an iXon+ EM CCD camera (Andor) and Micro-Manager software (37) (Edelstein et al., 2010). Observations were performed at room temperature (23°C). The images were processed by background subtraction and despeckle algorithms using the ImageJ software (National Institutes of Health, MD).

The central positions of the fluorescently-labeled dynein molecules were determined by the two dimensional (2D) Gaussian fitting of the image using Mark2 software (provided by Dr Kenya Furuta, National Institute of Information and Communications Technology, Kobe, Japan) (38) (Furuta and Toyoshima, 2008). Bright spots attached to microtubules longer than 0.5 s were analyzed. The diffusion constant (*D*) and drift velocity (*v*) of the dynein movements were estimated by calculating the mean square displacement (*ρ*) and fitting the following equation:

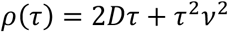

Where *τ* represents the non-overlapping time interval (39) (Qian et al., 1991). The mean run length and duration were calculated by fitting the data to cumulative distribution functions (11) (Imai et al., 2015).

### Microtubule gliding assay

Microtubule gliding assays were performed at 25°C. For the assays, glass chambers were sequentially coated with 1 mg/mL biotinamidocaproyl BSA (Sigma-Aldrich, St. Louis, MO), 1 mg/mL Neutravidin (Wako, Osaka, Japan), and 5 mg/mL α-casein (Merck, Darmstadt, Germany) (7) (Shima et al., 2006). A purified dynein-containing solution (including 50 nM dynein construct, 1 mM ATP, and 0.1 mg/mL casein) was then perfused into the chambers twice, at 5-min intervals. Finally, these chambers were filled with imaging solution containing 0.3 μM paclitaxel-stabilized non-labeled microtubules and then 1 mM ATP solution. MT gliding was observed under a BX-51 dark-field microscope (Olympus, Tokyo, Japan) with a 40× objective lens (NA 0.75; UPlanFl). The images were recorded using an sCMOS camera (Thorlabs) and analyzed using ImageJ.

### ATPase activity measurements

The microtubules were polymerized with 1 mM GTP for 30 min at 37°C and then stabilized with 20 μM paclitaxel (P-9600; LC Laboratories). Immediately before use, microtubules were spun down at 200,000 × *g* for 5 min using an Airfuge ultracentrifuge (Beckman-Coulter) to exclude free GTP. Basal and MT-activated ATPase activities were measured at 25°C using an EnzChek phosphate assay kit (E6646; ThermoFisher) (34) (Kon 2009) and a Tecan microplate reader (Infinite M200; Tecan).

### Microfluidics

To prepare a 0.1 × 0.1 × 25-mm flow chamber with portholes, polydimethylsiloxane (PDMS) was cast in a custom-built template (40) (Shirasaki et al. 2014), and then attached to the plasma-cleaned cover glass (No. 1; Matsunami). Briefly, the flow chamber was first washed with 50 μL of Milli-Q water. Then, 5% tetramethylrhodamin-labeled microtubules were immobilized on the glass surface a 0.1 mg/mL anti-tetramethylrhodamin antibody (ThermoFisher). It should be noted that the microtubules were placed in parallel owing to the shear flow. After blocking the surface with the imaging solution, HFB380-streptavidin coated polystyrene beads (diameter 200 nm; Polyscience) were introduced into the chamber and incubated for 3 min to allow binding with microtubules. The flow was driven by a syringe pump (YSP-201; YMC) using a 250-μL gastight syringe (41) (Hamilton; Shima et al., 2018): 0-10 μL/min flow of the imaging solution with 0.01% Methocel MC (Sigma-Aldrich) generated 0-6 pN of friction force on the beads. The flow rate was increased to 1 μL/min/s. The unbinding force (*F*) was calculated according to the Stokes-Einstein equation as follows:

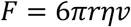

where *r, η*, and *v* represent the radius of the beads, viscosity, and flow velocity, respectively. Images of the beads and microtubules were recorded at a frame rate of 100 fps under the same setup for fluorescent microscopy. After the assay, the polarity of microtubules used here was determined by observing movement of DY-647-maleimide (Dyomics) labeled murine Kinesin-1 recombinant dimer (Shima et al., 2018; KIF5C, K395-His6-RCR) in the presence of 1 mM ATP. The kinesin construct was expressed in BL21-CodonPlus(DE3)-RIL (Agilent) and purified using TALON metal affinity resin (Clontech).

## Acknowledgments

We thank Rieko Shimo-Kon for her technical assistance. We also thank Karibu Sakai and Katsuhiko Minami for their help in the preparation of tubulin and maintenance of *Dictyostelium* stock. This work was supported by the Japan Society for the Promotion of Science KAKENHI (JP21J00021 to S.K.; 18K06147, 19H05379, and 21H00387 to T.S.; 20H05934 and 21H02441 to S.T.), the Ministry of Education, Culture, Sports, Science, and Technology (MEXT) grant JPMXP1020200101 to S.T., and the Japan Science and Technology Agency (JST) grant (JPMJCR1762) to S.T.

