## Supplementary material for "Key residue on cytoplasmic dynein for asymmetric unbinding and unidirectional movement along microtubule": Fig. S1-S2

**This PDF file includes:**

Figures S1-S2 and Table S1

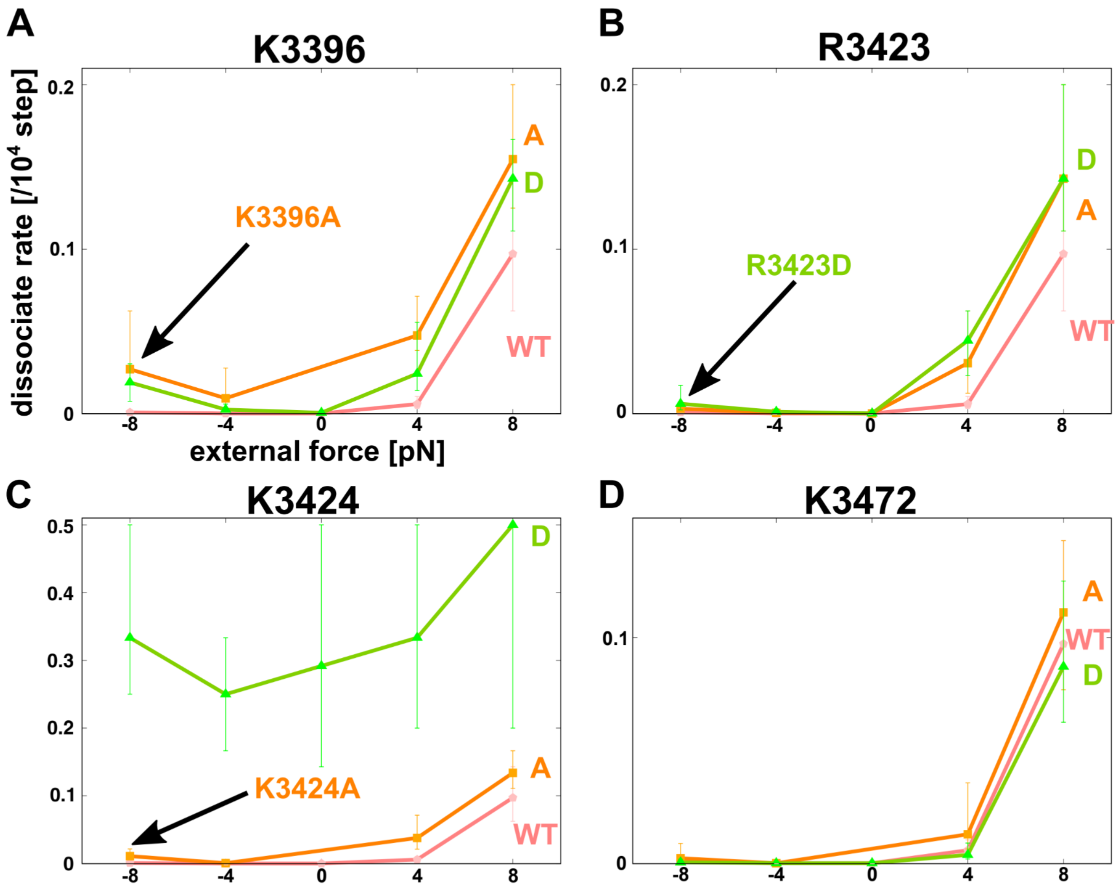


**Fig. S1. Effect of single point mutations of four surface residues in MTBD on simulated dissociate rates against various external force.**

The dissociate rate curves for the wild type MTBD (pink), alanine- (orange) and aspartate-substituted (green) mutants. The plots and error bars represent the mean values and standard deviations (SD), respectively; $N$ = 30 for each loading condition. The minus value of the force represents the loading toward the plus end of the MT. The target residues for the point mutations are K3396 (**A**), R3423 (**B**), K3424 (**C**) and K3472 (**D**).

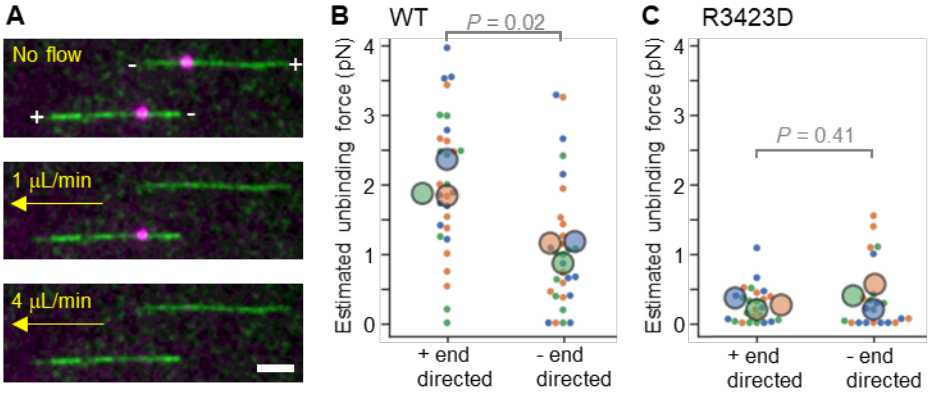


**Fig. S2. Unbinding force measurement using microfluidics.**

**(A)** Fluorescence images of surface-anchored microtubules (green) and dynein coated beads (magenta). The speed of flow (from right to left of the images) was increased from 0 to 10 µm/min. The flow is expected to apply up to 6 pN of friction force to the beads. After the force measurement, the polarity of microtubules was determined using fluorescently labeled Kinesin-1 (murine KIF5C, K411, ref. 41), a plus-end directed motor. Scale bar, 2 µm. (**B, C**) Dot plots of the force needed to unbind the beads from microtubules. The force was calculated from the flow rate when the beads detached from the microtubules. Each color represents a replicated experiment group. The big circles show the mean values of the groups. While the wild-type demonstrated asymmetry of the unbinding force (**B**, plus-end directed loading, 2.0 ± 1.0 pN, mean ± SD, n = 30; minus-end directed loading, 1.1 ± 0.9 pN, mean ± SD, n = 30; $P$ = 0.02, SuperPlots T-test), R3423 did not show significant difference upon loading directions (**C**, plus-end directed loading, 0.3 ± 0.3 pN, mean ± SD, n = 25; minus-end directed loading, 0.4 ± 0.5 pN, mean ± SD, n = 22; $P$ = 0.41, SuperPlots T-test).

| | | α-tubulin | | | | | | | | | | |
|---|---|---|---|---|---|---|---|---|---|---|---|---|
| | | V405 | H406 | W407 | Y408 | V409 | G410 | E411 | G412 | M413 | E414 | E415 |
| MTBD H1-H2 | P3400 | 0.0000 | 0.0000 | 0.0000 | 0.0000 | 0.0000 | 0.0000 | 0.0000 | -0.0349 | -0.0007 | -0.0432 | 0.0000 |
| | T3399 | 0.0000 | -0.0001 | 0.0000 | 0.0000 | 0.0000 | 0.0001 | 0.0000 | -0.0704 | -0.0674 | -0.0652 | 0.0002 |
| | P3398 | 0.0000 | -0.0001 | 0.0000 | -0.0424 | -0.0712 | -0.0706 | -0.0705 | -0.0700 | -0.0704 | -0.0705 | -0.0431 |
| | P3397 | 0.0000 | -0.0001 | 0.0000 | -0.0254 | -0.0716 | -0.0720 | -0.0718 | -0.0704 | -0.0712 | -0.0663 | 0.0004 |
| | K3396 | 0.0000 | -0.0566 | -0.0624 | -0.0722 | -0.0709 | -0.0707 | -0.0711 | -0.0702 | -0.0709 | 0.0012 | 0.0011 |
| | P3395 | 0.0000 | -0.0723 | -0.0737 | -0.0724 | -0.0710 | -0.0713 | -0.0725 | -0.0716 | -0.0018 | 0.0018 | 0.0014 |
| | L3394 | -0.0425 | -0.0730 | -0.0731 | -0.0726 | -0.0713 | -0.0720 | -0.0729 | -0.0713 | -0.0546 | -0.0066 | 0.0016 |
| | S3393 | -0.0327 | -0.0744 | -0.0201 | -0.0005 | -0.0726 | -0.0734 | 0.0003 | 0.0008 | 0.0006 | 0.0017 | 0.0015 |
| | K3392 | 0.0000 | -0.0093 | 0.0000 | 0.0000 | 0.0003 | -0.0004 | 0.0001 | 0.0003 | 0.0002 | 0.0003 | 0.0007 |
| | I3391 | 0.0000 | 0.0000 | 0.0000 | 0.0000 | 0.0001 | 0.0001 | 0.0000 | 0.0001 | 0.0000 | 0.0001 | 0.0002 |
| | E3390 | 0.0000 | -0.0004 | 0.0000 | 0.0000 | 0.0002 | 0.0001 | 0.0001 | 0.0003 | 0.0002 | 0.0010 | 0.0012 |

| | | β-tubulin | | | | | | | | | | |
|---|---|---|---|---|---|---|---|---|---|---|---|---|
| | | S153 | K154 | I155 | R156 | E157 | E158 | Y159 | P160 | D161 | R162 | I163 |
| MTBD H3 | K3425 | -0.0068 | -0.0010 | 0.0000 | -0.0082 | -0.0765 | 0.0001 | 0.0000 | 0.0000 | 0.0000 | 0.0000 | 0.0000 |
| | K3424 | -0.0760 | -0.0765 | -0.0649 | -0.0758 | -0.0737 | -0.0760 | -0.0415 | -0.0729 | 0.0000 | 0.0000 | 0.0000 |
| | R3423 | -0.0258 | -0.0124 | -0.0003 | -0.0754 | -0.0739 | -0.0734 | -0.0611 | -0.0753 | -0.0234 | 0.0000 | 0.0000 |
| | I3422 | 0.0000 | 0.0000 | 0.0000 | -0.0008 | -0.0757 | -0.0001 | 0.0001 | -0.0141 | 0.0000 | 0.0000 | 0.0000 |
| | D3421 | 0.0000 | -0.0001 | 0.0000 | -0.0262 | -0.0744 | -0.0733 | -0.0104 | -0.0329 | 0.0000 | 0.0000 | 0.0000 |
| | A3420 | -0.0001 | -0.0011 | -0.0033 | -0.0723 | -0.0728 | -0.0737 | -0.0742 | -0.0738 | -0.0675 | 0.0000 | 0.0000 |
| | W3419 | 0.0000 | 0.0000 | 0.0000 | 0.0000 | -0.0518 | -0.0117 | -0.0137 | -0.0703 | -0.0274 | 0.0000 | 0.0000 |
| | E3418 | 0.0000 | 0.0000 | 0.0000 | 0.0000 | 0.0001 | 0.0000 | 0.0000 | 0.0000 | 0.0000 | 0.0000 | 0.0000 |
| | L3417 | 0.0000 | 0.0000 | 0.0000 | 0.0000 | 0.0000 | 0.0000 | 0.0000 | 0.0000 | 0.0000 | 0.0000 | 0.0000 |
| | K3416 | 0.0000 | 0.0000 | 0.0000 | 0.0000 | 0.0000 | 0.0000 | 0.0000 | 0.0000 | 0.0000 | 0.0000 | 0.0000 |
| | K3415 | 0.0000 | 0.0000 | 0.0000 | 0.0000 | 0.0000 | 0.0000 | 0.0000 | 0.0000 | 0.0000 | 0.0000 | 0.0000 |

| | | α-tubulin | | | | | | | | | | |
|---|---|---|---|---|---|---|---|---|---|---|---|---|
| | | W407 | Y408 | V409 | G410 | E411 | G412 | M413 | E414 | E415 | G416 | E417 |
| MTBD H6 | G3475 | 0.0000 | 0.0000 | 0.0000 | 0.0000 | 0.0000 | 0.0000 | 0.0000 | 0.0000 | -0.0007 | 0.0000 | 0.0000 |
| | C3474 | 0.0000 | 0.0000 | -0.0001 | 0.0000 | 0.0000 | 0.0000 | 0.0000 | -0.0084 | -0.0141 | 0.0003 | 0.0001 |
| | A3473 | 0.0000 | 0.0000 | -0.0075 | 0.0000 | 0.0000 | 0.0003 | 0.0002 | -0.0028 | -0.0501 | 0.0017 | 0.0009 |
| | K3472 | 0.0000 | 0.0000 | -0.0041 | 0.0000 | 0.0000 | 0.0004 | 0.0009 | -0.0595 | -0.0700 | -0.0618 | 0.0018 |
| | S3471 | 0.0000 | 0.0000 | -0.0719 | -0.0001 | 0.0000 | -0.0201 | -0.0651 | -0.0695 | -0.0703 | -0.0692 | -0.0487 |
| | A3470 | 0.0000 | 0.0000 | -0.0370 | -0.0001 | 0.0000 | -0.0515 | -0.0693 | -0.0697 | -0.0711 | -0.0704 | -0.0700 |
| | R3469 | 0.0000 | 0.0000 | 0.0001 | 0.0001 | 0.0000 | 0.0007 | 0.0011 | -0.0702 | -0.0706 | -0.0701 | -0.0684 |
| | N3468 | 0.0000 | 0.0000 | 0.0000 | 0.0000 | 0.0000 | 0.0000 | 0.0000 | -0.0398 | -0.0690 | -0.0681 | 0.0000 |
| | V3467 | 0.0000 | 0.0000 | 0.0000 | 0.0000 | 0.0000 | 0.0000 | 0.0000 | -0.0494 | -0.0185 | -0.0436 | 0.0000 |
| | T3466 | 0.0000 | 0.0000 | 0.0000 | 0.0000 | 0.0000 | 0.0000 | 0.0000 | 0.0000 | 0.0000 | -0.0162 | 0.0001 |
| | E3465 | 0.0000 | 0.0000 | 0.0000 | 0.0000 | 0.0000 | 0.0001 | 0.0000 | 0.0000 | 0.0000 | 0.0001 | 0.0000 |

**Table S1. Contact probability of residue pairs between MTBD and MT.**

The difference values of contact probability between 4 pN of plus-end and minus-end directed loadings on each residue pair. The positively and negatively charged residues are colored in bule and red, respectively. The top 20 pairs in the difference values are highlighted in yellow.
